# Different modes of engagement with the nucleosome acidic patch yield distinct functional outcomes

**DOI:** 10.64898/2026.01.14.699602

**Authors:** Ujani Chakraborty, Emma Christina Saccone, Grisel Cruz-Becerra, Laiba F. Khan, Nina Arslanovic, Rhiannon Aguilar, Susan L. Gloor, Sabrina R. Hunt, Heather J. Folkwein, Natalia Ledo Husby, Keith E. Maier, Matthew R. Marunde, Noah K. Schomburg, Anup Vaidya, Martis W. Cowles, Bryan J. Venters, George Kassavetis, Zu-Wen Sun, James T. Kadonaga, Jean-Paul Armache, Michael-Christopher Keogh, Jessica K. Tyler

## Abstract

The nucleosome acidic patch is a hub of coordinated engagement by proteins that regulate genomic function. Here we report that *S. cerevisiae* Dot5 contains an arginine-rich HMGN-like motif that mediates nucleosome acidic patch binding and is required for the cell growth, DNA repair and heterochromatin defects exhibited when the protein is overexpressed. The heterologous expression of camelid single chain antibodies to the nucleosome acidic patch confers a similar range of phenotypes, with the most severe observed when an ‘arginine-anchor’ mode of binding analogous to many endogenous factors is employed. This highlights a delicate balance between nucleosome acidic patch interactors critical for normal cellular functions and dysregulated in disease.

## INTRODUCTION

Chromatin serves as a dynamic structure that packages the eukaryotic genome and regulates DNA-templated processes including replication, repair and transcription. The repeated structure is the nucleosome: ∼ 147 base pairs of DNA and a core histone octamer with two copies each of H2A, H2B, H3 and H4 (Kornberg 1974; Luger et al. 1997).

Epigenetic regulation is achieved by mechanisms that locally alter chromatin accessibility. This involves ATP-dependent remodeling, DNA methylation and histone post-translational modifications (PTMs), with each reliant on direct nucleosome interactions to control the recruitment and activity of regulatory and effector protein complexes (McGinty and Tan 2015; McGinty and Tan 2021; Peng et al. 2021a).

The core histones have unstructured and positively charged tails that engage the negatively charged DNA and ‘acidic patch’ of the same or an adjacent nucleosome, with this network of electrostatic interactions helping to compact DNA into chromatin (Hansen, Tse and Wolffe 1998; Horn and Peterson 2002; Gordon, Luger and Hansen 2005; Shogren-Knaak et al. 2006; Shogren-Knaak and Peterson 2006). In parallel, the various protein complexes that interact with nucleosomes often do so by engaging the local PTM landscape, wrapping and/or linker DNA, and the acidic patch. The last is thus of particular interest. It is formed by negatively charged residues from each histone H2A-H2B dimer (Luger et al.

1997), and provides a binding surface for the unacetylated H4 tail (Shogren-Knaak et al. 2006) and diverse ATP-dependent nucleosome remodelers, histone modifiers, DNA modifiers, DNA repair factors, transcription factors, cell cycle regulators, RNA metabolism proteins and chromatin architectural proteins (Skrajna et al. 2020; McGinty and Tan 2021; Oleinikov et al. 2023). Each typically binds the negatively charged patch via one or more positively charged arginines (Skrajna et al. 2020; Oleinikov et al. 2023), and such a recurrent mechanism raises a question: how is it coordinated and what would ensue from increased competition?

## RESULTS

### Yeast Dot5 binds to the nucleosome acidic patch via its HMGN-like motif *in vitro* and

#### in vivo

We recently showed that tardigrade (*Ramazzottius varieornatus*) Dsup contains a functional HMGN-like motif (RRSSRLTS) (Chavez et al. 2019; Aguilar et al. 2025; Alegrio-Louro et al. 2025) previously only described in vertebrates(McBryant, Adams and Hansen 2006). This arginine-rich motif (**Figure 1A**) binds the nucleosome acidic patch and is essential for Dsup to mediate its genoprotective role (Chavez et al. 2019; Aguilar et al. 2025; Alegrio-Louro et al. 2025). Budding yeast Dot5 also contains a HMGN-like motif (aa6-13, RRSARIAT: **Figure 1A** and **Supplemental Figure S1A**), encouraging us to examine any role in chromatin binding. To this end, we recombinantly expressed and purified Dot5 and a range of motif mutants (**Figure 1A** and **Supplemental Figure S1B**) and determined their binding to nucleosomes and free DNA by electrophoretic mobility shift. This identified more robust binding of the wild-type (WT) protein to nucleosomes over free DNA, with the nucleosome engagement lost on mutation (to alanine) or deletion of one or more motif arginines (R6, R7 or R10; **Figure 1C and Supplemental Figure S2**). This was further interrogated by Captify assay (**Supplemental Figure S3**), where Dot5 bound free DNA more tightly than nucleosomes at low salt (100 mM NaCl), but flipped this preference at an ionic strength closer to normal saline (150 mM; 0.9%) (**Figure 1D**). Deletion or mutation of the HMGN-like motif essentially ablated nucleosome binding but retained some engagement with free DNA (**Figures 1E-F**).

**Figure 1.**
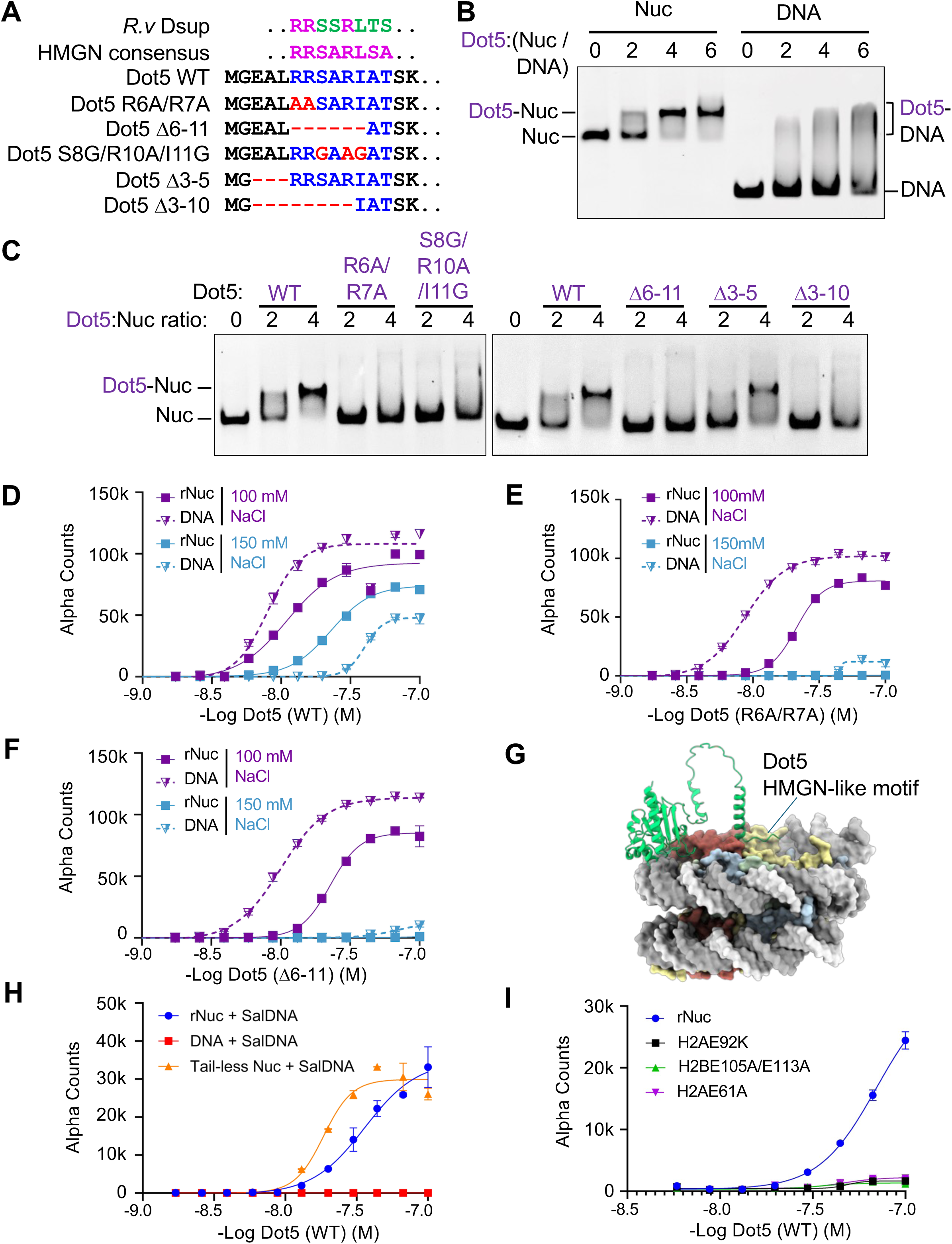
Yeast Dot5 preferentially binds via its HMGN-like motif to nucleosomes over free-DNA. **A.** *Saccharomyces cerevisiae* Dot5 contains a HMGN-like motif near its N-terminus (aa 6-13; RRSARIAT; see also **Supplemental Figure S1**), along with residues targeted in mutant alleles (*e.g.*, Dot5 D6-11). The related motif from *Ramazzottius varieornatus* Dsup (aa 363-370, RRSSRLTS) contains three functionally essential arginines. The vertebrate HMGN consensus sequence (RRSARLSA) is shown. **B.** Recombinant Dot5 (6His-Dot5-FLAG; **Supplemental Figure S1** and **Supplemental Table 1**) binds more effectively to mononucleosomes over free DNA. Reactants were combined as indicated and complexation analyzed by electrophoretic mobility shift assay (EMSA) on a nondenaturing 4.5% polyacrylamide gel. **C.** Conserved arginines (R6, R7 and R10) in the Dot5 HMGN-like motif are required for its association with nucleosomes (see also **Supplemental Figure S2**). Wild-type or mutant Dot5 protein was mixed with mononucleosomes as indicated, and complexation analyzed by EMSA on a nondenaturing 4.5% polyacrylamide gel. **D-F.** Impact of ionic strength (100 mM or 150 mM [normal saline, 0.9%]) on binding of Dot5 (WT) (**D**), (R6A, R7A) (**E**), or (D6-11) (**F**): 1 - 0 μM in 1.5-fold serial dilutions) to unmodified nucleosome (rNuc on 147 bp DNA; 10 nM) or free DNA (147 x 601: 2.5 nM). Interactions were assayed by Captify-Alpha (**Supplemental Figure S3**). **G.** AlphaFold3 model of Dot5-Nucleosome complexation predicts points of contact between the Dot5 (green) HMGN-like motif (labeled) and nucleosome acidic patch (histones H2A in yellow and H2B in brown), and between the Dot5 C-terminus and DNA. **H.** Salmon sperm DNA (salDNA; 1μg/ml) was a more effective competitor of Dot5 binding to free-DNA *vs*. nucleosome (rNuc or histone tails digested with trypsin [Tail-less]), suggesting multivalent engagement with the higher-order target. Reactions were in 150 mM NaCl + salDNA. **I.** Dot5 binding to nucleosomes is abolished by acidic-patch mutations. Reactions were in 150 mM NaCl + 1μg/ml salDNA.

Factors that engage nucleosomes most often contain multiple weak points of contact that synergize for effective binding (Peng et al. 2021b; Thomas et al. 2023; Weinzapfel et al. 2024; Aguilar et al. 2025; Keogh et al. 2025; Marunde et al. 2025). Indeed, AlphaFold3 (Abramson et al. 2024) modeling proposed Dot5 to bind the nucleosome at two positions: its N-terminal HMGN-like motif with the acidic patch, and a region of its C-terminus with DNA (**Figure 1G**). We explored and supported this further by adding salmon sperm DNA (salDNA) to Captify assays (Dilworth et al. 2022; Marunde et al. 2022; Aguilar et al. 2025; Marunde et al. 2025), which competed the binding to free DNA but left that to nucleosomes (**Figure 1H**). Suggesting that this engagement is synergistic, under these assay conditions Dot5 showed even stronger binding to nucleosomes lacking histone tails (**Figure 1H**); likely because of increased accessibility to the nucleosomal DNA usually associated with these tails (Skrajna et al. 2020). Finally, under the same competitive conditions, Dot5 had profoundly reduced binding to nucleosomes with acidic patch mutations (([H2AE61A]_2_), ([H2AE92K]_2_) or ([H2BE105A/E113A]_2_); **Figure 1I**), confirming this as a primary point of engagement. Given the above, we predicted Dot5 would be associated with *in vivo* chromatin, and this was confirmed after cell fractionating and immunoblotting a yeast strain expressing Dot5-FLAG from its endogenous locus (**Supplemental Figure S4A**). However, this binding was not targeted to any genomic locations during yeast exponential growth, since CUT&RUN failed to identify any regions of Dot5-FLAG enrichment (**Supplemental Figure S4B**).

### Binding of the Dot5 HMGN-like motif or VHH to the nucleosome acidic patch disrupts growth, silencing and genome integrity

A major strength of yeast-based studies is the ease of exploring cellular phenotypes in the context of factor deletions, mutations or altered expression levels (Hampsey 1997).

We first deleted Dot5 but failed to reproduce the sensitivity to diverse genotoxins (Cha et al. 2003; Wong, Siu and Jin 2004) in either a W303 or S288C yeast strain background (**Supplemental Figure S5**). We thus turned to over-expression, since Dot5 had previously been identified to disrupt heterochromatin function under such conditions; the origin of its name (Disrupter of Telomeric Silencing 5) (Singer et al. 1998). Dot5-FLAG was expressed from the *GAL1* promoter, maintained in yeast on a high-copy 2μ plasmid, and expression confirmed titratable by galactose (Mumberg, Muller and Funk 1994) (**Supplemental Figure S6A**). Under these conditions Dot5 (R6A/R7A)-FLAG was unstable but Dot5 (Δ6-11)-FLAG effectively expressed (**Figure 2A**). However, the latter did not associate with the chromatin fraction (**Figure 2B**) unless highly expressed (**Supplemental Figure S6B**), confirming the importance of the Dot5 HMGN-like motif for nucleosome binding (**Figure 1**). When overexpressed, Dot5 (WT) compromised cell growth and reporter gene repression at the telomere (*TELVIIL::URA3*) and silent mating locus (*HMR::ADE2*) (**Figure 2C-E**). However, yeast overexpressing Dot5 (Δ6-11) displayed much more subtle versions of these phenotypes (**Figure 2C-**E **and S**u**pplemental Figure S6C**), suggesting they manifest when the protein is robustly engaged with chromatin.

**Figure 2.**
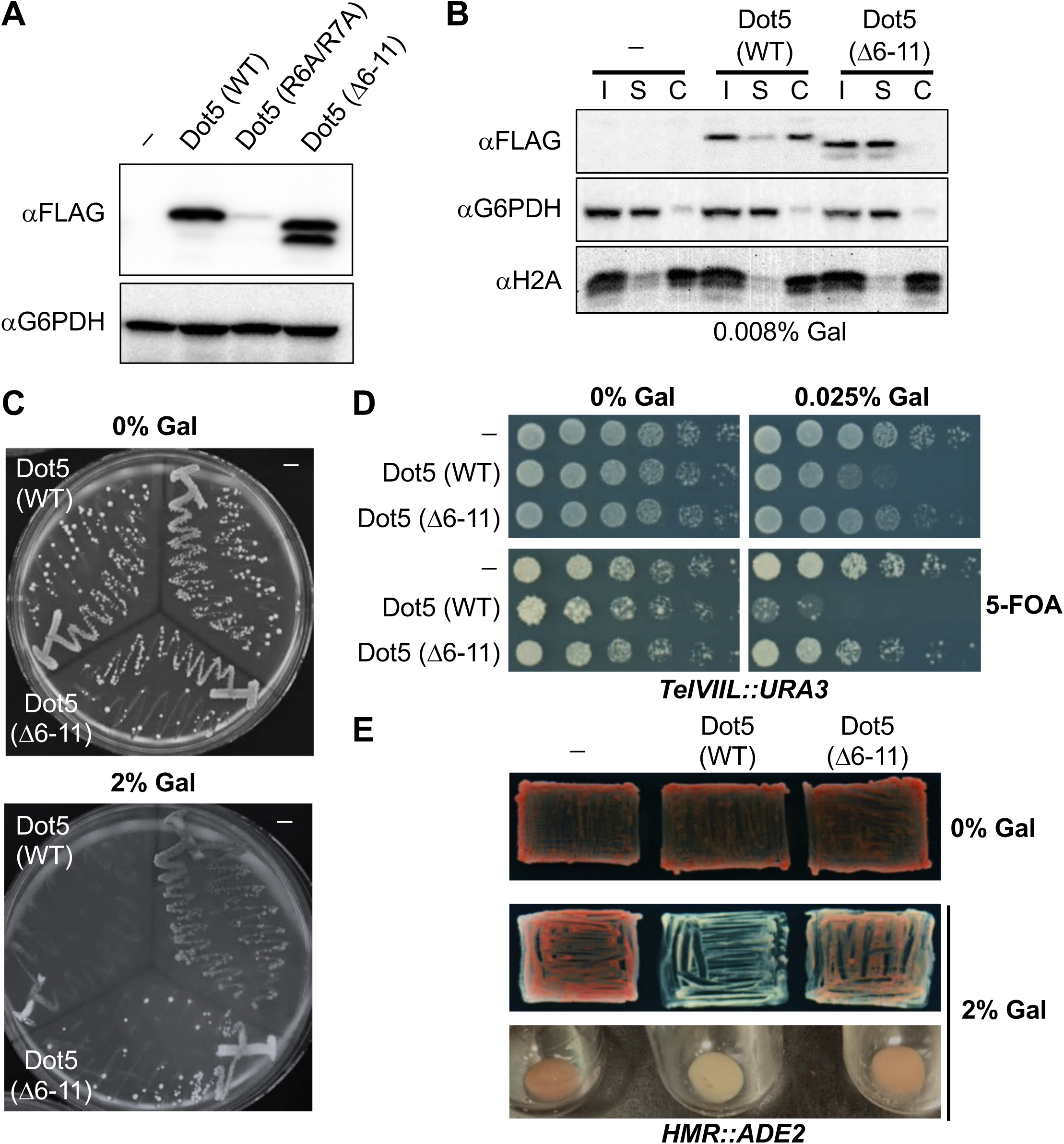
Heterologous expression and chromatin binding of Dot5 inhibits yeast growth and heterochromatin function. **A.** Galactose-induced expression of Dot5-FLAG alleles were assayed by immunoblot. G6PDH is a loading control for each protein extract. “-” refers to yeast containing empty vector. **B.** Dot5 (WT) associates with chromatin more effectively than HMGN-mutant Dot5 (D6-11). Expression of each FLAG-tagged allele was titrated by galactose addition (see also **Supplemental Figure S6**), before yeast cultures were spheroplasted (Input), resolved to <u>S</u>oluble and <u>C</u>hromatin fractions, and immunoblotted. Confirming effective fractionation: G6PDH is a soluble protein, H2A is chromatin bound. **C.** Overexpression of Dot5 (WT) but not Dot5 (D6-11) compromises cell growth. Yeast (ROY1172) harboring *2μ* galactose inducible plasmids expressing the indicated proteins (**Supplemental Table 1**) were struck on SC leucine dropout (SC-leu) plates supplemented with 2% raffinose-/+ 2% galactose. SC, synthetic complete. **D.** Overexpression of Dot5 (WT) but not Dot5 (D6-11) disrupts telomeric silencing. Yeast containing a *TelVIIL::URA3* reporter were spotted on SC-leu plates-/+ 5-Fluoroorotic acid (5-FOA). Derepression of sub-telomeric *URA3* allows the conversion of 5-FOA to toxic 5-fluorouracil, leading to cell death. **E.** Overexpression of Dot5 (WT) but not Dot5 (D6-11) compromises HMR silencing. Yeast containing an *HMR::ADE2* reporter were patched on SC-leu plates (or cultured in similar liquid media) supplemented with 2% raffinose-/+ 2% galactose. Red color indicates effective silencing of the *ADE2* reporter; white color its derepression.

The nucleosome acidic patch is a major hub of engagement for diverse chromatin regulators (Skrajna et al. 2020; Lagadec, Parissi and Lesbats 2022; Wesley et al. 2022; Oleinikov et al. 2023), suggesting the need for some degree of coordination to prevent pathological interference. We surmised that the phenotypes of overexpressed Dot5 (WT) were due to using its HMGN-like motif to compete for acidic-patch occupancy. To explore this further we attempted to express in yeast a range of nucleosome acidic-patch binders as potential direct competitors; each with a Nuclear Localization Signal and 3xHA epitope tag (**Table S1**). Here the Dot5 N-terminus (residues 1-20) and Kaposi’s Sarcoma Herpesvirus LANA(Barbera et al. 2006) (residues 1-22; previously used to *in vivo* tether or *in vitro* compete interactions with the nucleosome acidic patch (Wilson et al. 2016; Nozaki and Kanai 2021)) were likely unstable (**Supplemental Figure S7**) and thus removed from further consideration. However, Dot5 (residues 1-56) was effectively expressed, as were three camelid single-chain antibodies (also termed VHH; Variable Heavy domain of a Heavy chain) to the nucleosome acidic patch: Chromatibody (Cb), 1B2 and 1G1 (Jullien et al. 2016; Hicks et al. 2024; Marunde et al. 2025) (**Figure 3A**). All four of these entities showed some association with chromatin (**Figure 3B**), and all four enhanced the sensitivity of WT and *rad52*Δ yeast to phleomycin and zeocin, a genome integrity defect shared with strains overexpressing Dot5 (WT) (**Figure 3C** and **Supplemental Figure S8A**). In further phenotypic testing, overexpression of Dot5 (WT) and VHH 1B2 led to profound growth and silencing defects, compared to the weaker defects of VHH Cb and Dot5 (1-56), and negligible impact of VHH 1G1 (**Figure 3D-E** and **Supplemental Figure S8B-C**). We surmised that the weaker overexpression phenotypes of Dot5 (1-56) relative to Dot5 (WT) were because the former does not include the DNA binding region of the full-length protein (**Figure 1G**) and is thus less effectively associated with chromatin (compare **Figures 3B** and **2B**).

**Figure 3.**
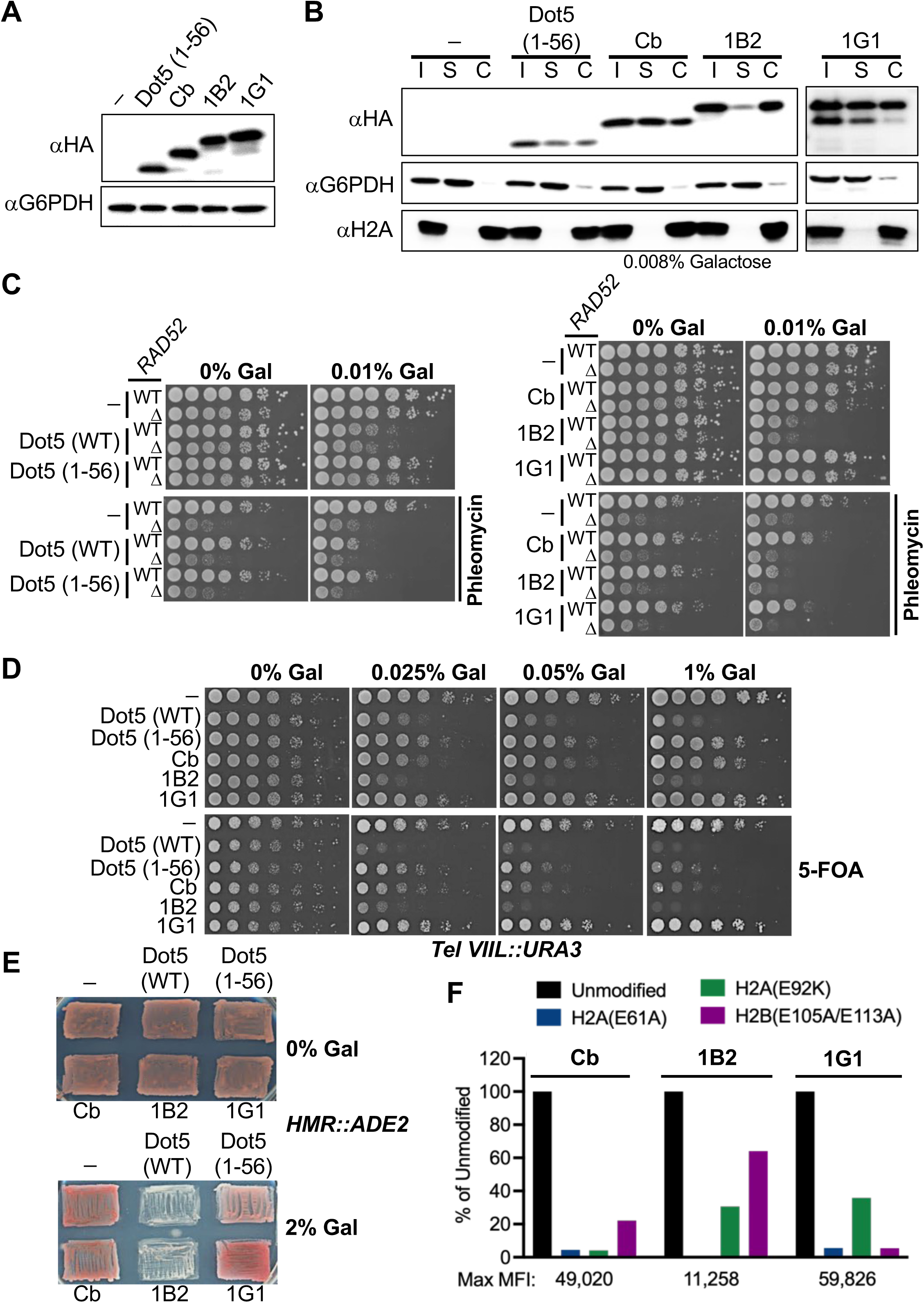
Expression of nucleosome acidic patch binding Variable Heavy domain of a Heavy chain (VHH) compromises yeast cell growth, DNA repair and/or heterochromatin function. **A.** Galactose-induced expression of Dot5 (1-56) containing the HMGN-like motif (aa6-13) or nucleosome acidic-patch binding VHH (Cb, 1B2 and 1G1) (**Supplemental Table 1**) was assayed by immunoblot. G6PDH is a loading control for each protein extract. “-” refers to yeast containing the empty vector. Cb, chromatibody. **B.** Heterologous proteins show varying degrees of chromatin association. Yeast cultures were spheroplasted (Input), resolved to <u>S</u>oluble and <u>C</u>hromatin fractions, and immunoblotted. Confirming effective fractionation: G6PDH is a soluble protein, H2A is chromatin bound. **C.** Influence of expression of Dot5 (WT), Dot5 (1-56) and VHH (Cb, 1B2 and 1G1) on cell growth (see also **Supplemental Figure S8A, S8C**) and DNA repair. Strains (*RAD52* or *rad52*D background) harboring *2μ* galactose inducible plasmids expressing the indicated proteins were spotted on SC-leu plates supplemented with 2% raffinose, galactose as indicated,-/+ phleomycin (10 mg/ml) as a DNA damaging agent. **D.** Influence of expression of Dot5 (WT), Dot5 (1-56) and VHH (Cb, 1B2 and 1G1) on telomere-proximal silencing, as described in legend to **Figures 2D**-**E**. Overexpression of Dot5 (WT) and VHH 1B2 similarly compromise HMR silencing (see also **Supplemental Figure S8B**), as described in legend to **Figures 2E-F**. Expression of each VHH (Cb, 1B2 and 1G1) yields differential sensitivity to nucleosome acidic patch mutants (H2AE61A, H2AE92K or H2BE105A/E113A) in Captify-Luminex assays (see Methods). Data is presented as % binding relative to unmodified nucleosome. Max MFI, maximum Median Fluorescence Intensity.

### Differential recognition of the nucleosome acidic patch by VHH 1B2 and 1G1

In prior studies (Hicks et al. 2024) we measured the binding of each VHH (Cb, 1B2 and 1G1) to nucleosomes at 559 nM, >2 μM and >2 μM respectively; which for comparison was weaker than GST-LANA at 192 nM (all expressed as EC_50_^Rel^). Intrigued by the very different growth and silencing yeast phenotypes resulting from expressing each VHH, we set out to further delineate their engagement with the nucleosome acidic patch, and observed differential sensitivity to the H2AE61A, H2AE92K or H2BE105A/E113A mutants (**Figure 3F**). For further insight, we sought to structurally define the nucleosomal interactions of 1B2 and 1G1 as of particular interest, given their similar affinity for nucleosomes (Hicks et al. 2024) but representing each extreme of *in vivo* phenotype. Each VHH was mixed 10:1 with mononucleosome (on 147bp 601 DNA) and prepared for cryo-EM without fixation (see Methods). Both yielded high-resolution structures (1B2-Nucleosome at 3.17 Å; 1G1-Nucleosome at 3.10 Å) suitable to map protein folding and localize side-chains, particularly at the interaction interface **(Figure 4A** and **Supplemental Figures S9-S11**). Camelid VHH paratope diversity arises from a combination of differential germline gene usage and somatic hypermutation to generate highly diverse and structurally flexible complementarity determining regions (CDR1-3) and often also involvement of the intervening framework regions (FW1-3) (Nguyen et al. 2000; Mitchell and Colwell 2018; Wang et al. 2024). In the case of 1B2 and 1G1, their acidic-patch binding paratopes were respectively distributed across two or four points of contact (FW1 and CDR3; FW1/CDR1, CDR2, FW2, and CDR3), with 1G1 having a more discontinuous paratope (**Figure 4B-C** and **Supplemental Figure S12**). This could suggest a requirement for greater co-ordination by 1G1 to achieve effective target binding.

**Figure 4.**
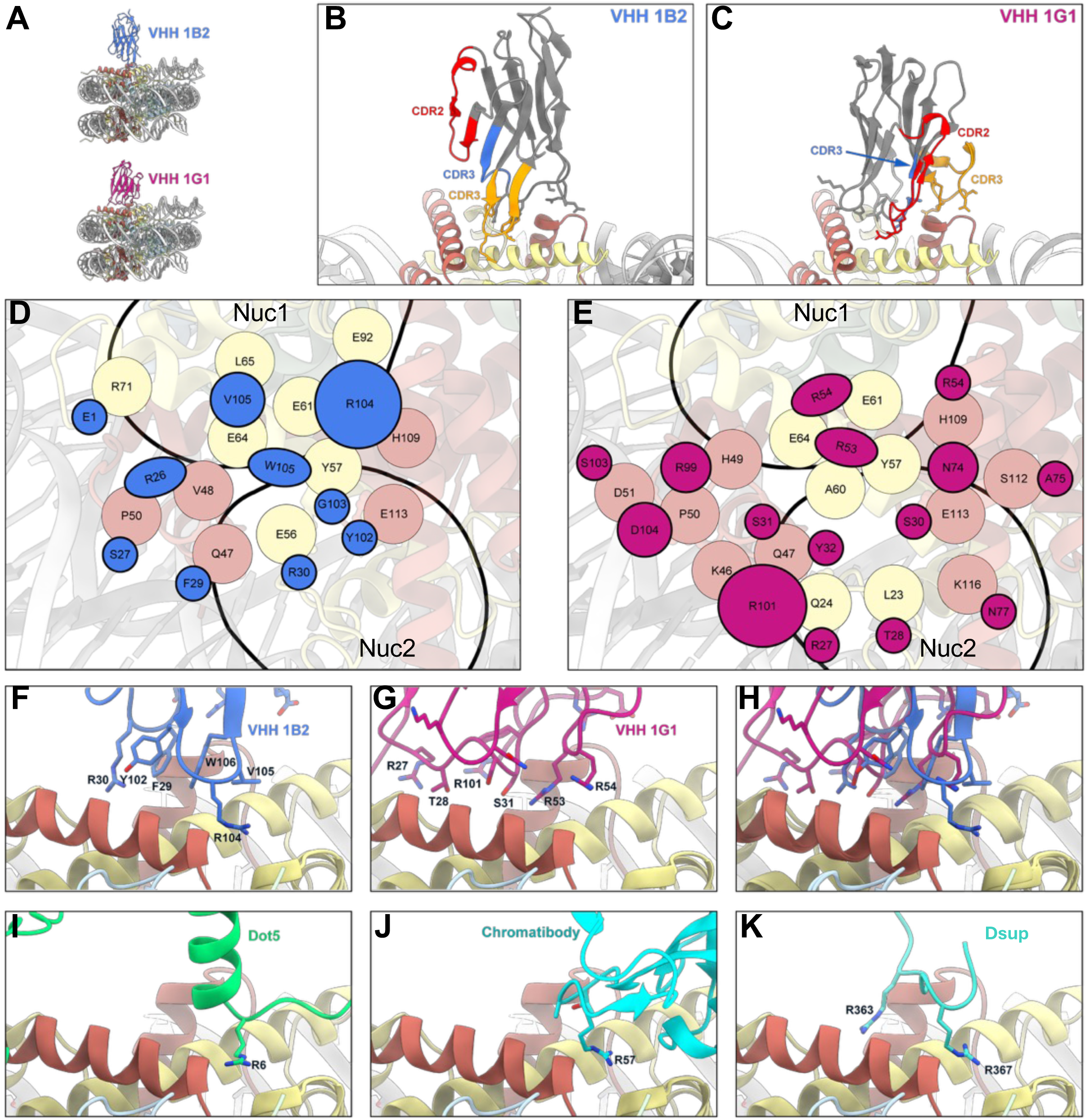
Cryo-EM reveals distinct modes of nucleosome acidic patch binding by VHHs 1B2 and 1G1. **A.** Atomic models of VHH 1B2 (blue at 3.17 Å, top) and 1G1 (pink at 3.10 Å, bottom) bound to the nucleosome; each built from their respective cryo-EM reconstructions. **B-C.** 1B2 and 1G1 are colored by their FW-CDR domains (primary sequence of each paratope in **Supplemental Figure S12**). **D-E.** Residues of the nucleosome acidic patch (top-down view) engaged by 1B2 (**D**, blue) and 1G1 (**E**, dark pink). Circle size for 1B2 and 1G1 residues is proportional to their contacts with residues from histone H2A (yellow) and H2B (salmon pink). Nuc1, acidic patch. **F-K.** Comparison of the mode of binding of 1B2 (**F**, Arginine anchor at R104; PDB 9ZEN), 1G1 (**G**, PDB 9ZEO), 1B2 *vs* 1G1 (**H**), Dot5 (**I**, AlphaFold3 model), Cb (**J**, AlphaFold3 model) and Dsup (**K**, PDB 9D3L).

As predicted from their mutant sensitivity profiles (**Figure 3D-F**), each VHH engaged a different area of the acidic patch (**Figure 4D-E**). Here 1G1 had a distributed interaction, including with the ‘Nuc2’ region (Skrajna et al. 2020; Oleinikov et al. 2023) (**Figure 4D-H**). In contrast, 1B2 occluded a larger area of ‘Nuc1’ (the central acidic patch) and used R104 as a main point of nucleosome contact (**Figure 4D-H**). This is reminiscent of the ‘arginine anchor’ employed by the HMGN-like motif (Alegrio-Louro et al. 2025), the Dot5 HMGN-like motif (**Figure 4I**), Cb (**Figure 4J**), Dsup (Alegrio-Louro et al. 2025) (**Figure 4K**) and factors from multiple biological pathways to engage the acidic patch (Barbera et al. 2006; Armache et al. 2011; Kato et al. 2011; Kato et al. 2013; McGinty, Henrici and Tan 2014; McGinty and Tan 2016) (**Supplemental Figure S13**). Overexpression of such entities would be presumed particularly competitive with the regular *in vivo* interplay of nucleosome interactors (**Supplemental Figure S14**).

## DISCUSSION

In this study we demonstrate the impact of heterologous competition for the nucleosome acidic patch, a hub of engagement for factors regulating diverse cellular functions. The overexpression of *Sc* Dot5, which uses an ‘arginine anchor’ in its HMGN-like motif to mediate acidic patch binding (**Figure 4I**), was profoundly disruptive to yeast cell growth and gene silencing (**Figures 2** and **3C**). These defects were shared with yeast expressing camelid VHH 1B2 but absent from those expressing VHH 1G1 (**Figure 3**). Each of these VHH binds nucleosomes with similar affinities (Hicks et al. 2024), but at different areas of the acidic patch, via distinct modes of engagement (respectively coordinating two or four paratope surfaces), and with 1B2 using an ‘arginine anchor’ mechanism (**Figures 3F** and **4**). As such, mimicking the means of acidic patch binding used by endogenous factors from multiple biological pathways (Barbera et al. 2006; Armache et al. 2011; Kato et al. 2011; Kato et al. 2013; McGinty, Henrici and Tan 2014; McGinty and Tan 2016) can prove highly disruptive to cellular function (**Supplemental Figure S14**).

The negatively charged acidic patch is engaged by the histone H4 tail (via positively charged residues 16-23) to promote higher order structure (Shogren-Knaak et al. 2006), and a diversity of proteins to recruit and regulate their activity. Indeed, competition between these many elements is integral to biology, with H4K16ac neutralizing the histone tail charge to relax inter-nucleosomal interactions (Kan, Caterino and Hayes 2009; Sinha and Shogren-Knaak 2010), while the binding of SOX pioneer transcription factors displaces the H4 tail to destabilize chromatin packing and create a transcriptionally permissive environment (Dodonova et al. 2020). Further, the mutation or modified abundance of acidic-patch binding proteins can lead to developmental and immune defects, intellectual disabilities, neurological disorders and cancer (Kadoch et al. 2013; Valencia et al. 2019; McBride et al. 2020; Nanduri, Furusawa and Bustin 2020; Lagadec, Parissi and Lesbats 2022). It is tempting to speculate that some of these phenotypes are directly due to perturbating the usual interplay on the nucleosome acidic patch.

Dot5 overexpression compromises gene silencing (Singer et al. 1998), DNA repair and cell growth when engaged with chromatin via its N-terminal HMGN-like motif and C-terminal DNA binding element (**Figures 1, 2** and **3C**). The arginine-rich acidic patch-binding HMGN motif was originally only identified in vertebrates, until our recent demonstration of a similar element in tardigrade Dsup (Aguilar et al. 2025; Alegrio-Louro et al. 2025). That protein confers a resistance to oxidative DNA damage on expressing cells; a genoprotection that requires effective chromatin binding. This is mediated by the multivalent engagement of Dsup with various nucleosome surfaces, including the wrapping DNA, histone tails and acidic patch (Aguilar et al. 2025; Alegrio-Louro et al. 2025). So why would the heterologous expression of one HMGN-like motif containing protein (Dot5) compromise cellular function, while another (Dsup) is protective? There are multiple variables. It may be simple abundance: Dsup has no impact at low levels, is protective at a histone-like stoichiometry, but codon-optimization to express at still higher levels yields inviable yeast (Aguilar et al. 2025) and neuronal toxicity (Escarcega et al. 2023), suggesting too much is deleterious. Alternatively, Dsup-nucleosome engagement is highly dynamic (Aguilar et al. 2025; Alegrio-Louro et al. 2025), which would be predicted for a protein that can ‘coat’ chromatin while being minimally disruptive to ongoing transactions such as DNA replication and transcription. Moreover, Dsup-nucleosome engagement is impressively multivalent, where interaction of its HMGN-like motif with the acidic patch or C-terminal adjacent sequences with DNA are largely redundant to mediate chromatin binding and genoprotection (Aguilar et al. 2025).

However, Dsup is not entirely without detriment on normal cellular function. Yeast expressing Dsup at a genoprotective level concurrently show elevated expression of genes proximal to telomeres or encoded from Ty transposable elements (Aguilar et al. 2025). Both are usually repressed locations (Curcio, Lutz and Lesage 2015), suggesting some opening of silent chromatin, reminiscent of the Dot phenotype.

Reduced growth and dysregulated silencing are commonly (but not universally) observed when over-expressing proteins that contain nucleosome acidic patch binding entities (**Supplemental Table 4**). However, this would not rise to a definitive demonstration of acidic patch competition, since each of these proteins contains a range of functional domains. However, our expression of mono-functional acidic patch binding VHH has allowed us to isolate ‘competition with endogenous interactors’ as a potential mechanism. It is not possible to identify any single factor impacted, and indeed it may be multi-factorial: the essential Epl1, Orc1, Spt16 and Sth1 are candidates for inviability; Sir3 for silencing defects; Arp5, Bre1, Ies2, Sgf11, Snf5 and Spt16 for genotoxin sensitivity (**Supplemental Table 4**).

It is likely that interfering with the normal landscape of acidic patch binding to a pathological degree requires a conflation of circumstances, including competitor abundance and means of engagement (how, where, and with what stability). Many endogenous factors use an ‘arginine anchor’, as does the HMGN (Alegrio-Louro et al. 2025), the HMGN-like motif of Dsup (Alegrio-Louro et al. 2025) (**Figure 4K**), and likely also the HMGN-like motif of Dot5 (**Figure 4I**) and the paratope of VHH Cb (**Figure 4J**). Here truncated Dot5 (1-56) and Cb confer one or more moderate phenotypes as their expression level rises. However, the impact of Dot5 (WT) is more severe, most likely because the full-length protein has an additional DNA-binding element (**Figure 1G**) that synergizes to stabilize its nucleosome engagement (**Figures 1D-F** and **2B**) and increase its disruptive potential (**Figure 2C-E**). We note that VHH 1B2 binds the nucleosome acidic patch using only two points of paratope - epitope engagement (thus easy to co-ordinate; indeed, observed as highly stable in cryo-EM) and including an ‘arginine anchor’ (**Figure 4** and **Supplemental Figure S12**). This would be imagined as the most effective strategy to compete myriad endogenous factors, and indeed it is the VHH with greatest phenotypic impact (**Figure 3**).

Highlighting its disease relevance, the nucleosome acidic patch is engaged by many viral proteins (Reinhardt et al. 2005; Barbera et al. 2006; Delelis et al. 2010; Aydin and Schelhaas 2016; Fang et al. 2016; Lagadec, Parissi and Lesbats 2022), and its residues are mutated in a range of ‘oncohistones’ (Nacev et al. 2019). These include H2AE56, H2AE61, H2AD90, H2AE92 and H2BE113; indeed, the last was the fifth most common site of mutation in one large scale oncohistone study (Nacev et al. 2019). It has been speculated that these mutations disrupt the association / function of effector complexes, leading to oncogenesis. We propose that the same impact could be mediated by the aberrant expression of proteins with nucleosome acidic patch binding domains, leading to a changed competitive landscape, and deleterious phenotypes such as genome instability and transcriptional dysregulation. This would also suggest that extreme care be taken when engaging the nucleosome acidic patch with experimental or therapeutic intent, as with binding peptide fusions (Chodaparambil et al. 2007; Dos Santos Passos et al. 2021), Dsup (Kirtane et al. 2025), or small molecules (Davey et al. 2017).

## MATERIALS AND METHODS

### Yeast strains, primers and plasmids

Yeast strains and plasmids (**Supplemental Table 1**) were created and maintained using standard methods (Rose 1990). *DOT5* was deleted in the S288c and W303 backgrounds by PCR amplifying *dot5Δ*::*KANMX* from the yeast knockout collection (Brachmann et al. 1998) and transformation using standard procedures (Gietz and Schiestl 1991).

Strains expressing Dot5-FLAG (strains UCY003 and UCY004) were made by CRISPR-Cas9 mediated genome editing (Aguilar, Shen and Tyler 2022) (guide RNAs and homologous template sequences in **Supplemental Table 1**). UCY005 (expressing Dot5 (Δ6-11)-FLAG) was made by CRISPR-Cas9 mediated gene editing of UCY003.

To construct Dot5 yeast expression plasmids (**Supplemental Table 1**), *DOT5-FLAG* was PCR amplified from UCY003 and cloned to the SalI and BamHI sites of p425-GAL1. Similarly, the *dot5-(Δ6-11)-FLAG* insert was PCR amplified from UCY005. p425-GAL1 containing codon-optimized Dot5 [1-56]-2xNLS-3xHA was synthesized by *Synbio technologies*. To construct VHH (Cb, 1G1 or 1B2) yeast expression plasmids (**Supplemental Table 1**), each was synthesized as codon-optimized in format VHH-2xNLS-3xHA (2x SV40 large tumor antigen NLS (PKKKRKV<u>PKKKRKV</u>)(Aguilar et al. 2025) and C-terminal 3xHA tag (3x YPYDVPDYA)) and cloned to the SalI and BamHI sites of p425-GAL1. All galactose titratable expression plasmids were transformed and tested in the ROY1172 background.

### Expression and purification of recombinant Dot5

The cloning, expression, and purification of wild-type and mutant versions of Dot5 closely followed the non-denaturing expression and purification of Dsup (Chavez et al. 2019). Briefly, the *E. coli* codon-optimized cDNA of Dot5 was synthesized with a C-terminal FLAG-tag (Gblock, *IDT*), inserted as an NcoI-XhoI fragment into pET21b-His6-Nco, and analyzed by DNA sequencing. The five mutant Dot5 sequences were obtained by PCR with upstream primers (starting 6 bp upstream of the NcoI site and containing the desired mutations) and the T7 terminator primer downstream for insertion as an NcoI-XhoI fragment into pET21b-His6-Nco. The expression of each plasmid in *E. coli* Rosetta (DE3) pLysS, with lysis, sonication and sequential chromatography on Ni-NTA agarose and anti-FLAG M2 agarose as previously (Chavez et al. 2019).

### Electrophoretic mobility shift analysis (EMSA)

147-bp 5S rDNA mononucleosomes were reconstituted essentially as previously (Chavez et al. 2019), except they contained HeLa core histones. Mononucleosomes (66 nM final) or free DNA (66 nM final) were combined with purified Dot5 (WT or mutant; amounts as indicated) in HEG buffer (25 mM HEPES (K^+^) pH 7.6, 0.1 mM EDTA, 10% (v/v) glycerol) containing 96 mM KCl, 0.01% NP-40 and 93 ng BSA in a total volume of 15 µl, incubated at 27 °C for 40 mins, and then loaded on a nondenaturing 4.5% polyacrylamide gel that had been pre-equilibrated at 4 °C and pre-run at 3.5 V/cm for 30 mins. Gels were run at an additional 4.5 hr at 3.5 V/cm and 4 °C, followed by ethidium bromide staining for visualization.

### Chromatin fractionation analysis

Chromatin fractionation was performed as previously (Aguilar et al. 2025). After spheroplasting, half of the lysate was removed (Input) and the other half centrifuged (16,000 g for 15 mins at 4 °C) for separation to supernatant (Soluble fraction) and pellet (Chromatin fraction). All three samples were made to 1x with 5x Laemmli buffer, boiled for 5 mins, clarified by centrifugation, 7.5% of the total volume resolved by 12.5% SDS-PAGE, membrane transferred, and immunoblotted with anti-FLAG (*Sigma* F1804; 1:1,000) to detect Dot5, anti-HA (*Biolegend* 901515; 1:750) to detect the Dot5 (1-56), Cb, 1G1 and 1B2, anti-H2A (*Active Motif* 39235; 1:2,000) as a chromatin bound protein, and anti-G6PDH (*Sigma* A9521; 1:25,000) as a non-chromatin bound protein (all antibodies are listed in **Supplemental Table 1**).

### Immunoblot analysis

Immunoblotting was performed as previously(Aguilar et al. 2025) using standard approaches. Cell pellets from mid-log cultures were collected by centrifugation, resuspended in lysis buffer (150 mM NaCl, 25 mM Tris pH 8.0, 1 mM EDTA, 10 mM β-mercaptoethanol, 1 mM PMSF), and mechanically lysed by votexing with zirconia beads. Clarified lysates were normalized for total protein by Bradford assay, resolved by reducing 12.5% SDS-PAGE, transferred to a nitrocellulose membrane (*Amersham Protran* 10600002), and blocked for 1 hr at room temperature. Immunoblotting was by overnight incubation at 4 °C with primary antibody (**Supplemental Table 1**): anti-FLAG (*Sigma* F1804; 1:1,000), anti-H2A (*Active Motif* 39235; 1:2,000), anti-G6PDH (*Sigma* A9521; 1:25,000), or anti-HA (*Biolegend* 901515; 1:750). Membranes were washed, incubated (1 hr at room temperature) with horseradish peroxidase-conjugated goat anti-mouse (*Promega* W4021; 1:2,500) or goat anti-rabbit (*Promega* W4011; 1:5,000) and washed again. HRP signal was detected using ECL Western Blotting Detection Reagent (*Cytiva Amersham*) and imaged using the *Proteinsimple* Fluor Chem E gel doc.

### Serial dilution spotting assays

For testing sensitivity to oxidizing agents, single colonies were inoculated in 5 ml YPD (WT, *dot5*Δ) or synthetic complete leucine dropout (SC-leu) media supplemented with 2% raffinose (strains containing overexpression plasmids), grown to saturation at 30 °C, made to OD_600_ = 1, and serially diluted (10-fold or 5-fold increments as indicated) in sterile distilled water. Spots were on either YPD plates or SC-leu 2% raffinose, galactose plates containing the indicated oxidizing / DNA damaging agents. Plates were incubated at 30 °C for 4-5 days before imaging.

For the survival assay, single colonies were inoculated in 5 ml SC-leu media supplemented with 2% raffinose, grown to saturation at 30 °C, made to OD_600_ = 1, and serially diluted (5-fold) in sterile distilled water. Spots were on SC-leu plates (2% raffinose, galactose as indicated). Plates were incubated at 30 °C for 4-5 days before imaging.

For telomeric silencing assays (using the *TELVIIL∷URA3* reporter: **Supplemental Table 1**), single colonies were inoculated in 5 ml SC-leu media with 2% raffinose as carbon source, grown to saturation at 30 °C, made to OD_600_ = 1, and serially diluted (5-fold increments in sterile distilled water). Spots were on SC-leu plates (2% raffinose; galactose and-/+ 1 mg/ml 5-Fluoroorotic acid (5-FOA) as indicated). Plates were incubated at 30 °C for 4-5 days before imaging.

### Mating type locus silencing assays

To assay heterochromatin function at the silent mating loci (using the *HMR∷ADE2* reporter: **Supplemental Table 1**), single colonies from SC-leu plates containing 2% raffinose as carbon source were either: (i) patched on SC-leu plates (supplemented with 2% glucose or 2% galactose), incubated at 30 °C, and imaged after three to four days; or (ii) grown in SC-leu media containing 2% galactose for eight days, equal OD_600_ by centrifugation, and pellet color assessed.

### Recombinant nucleosomes (rNuc)

All mononucleosomes (*EpiCypher*; **Supplemental Table 1**) were created from fully-defined (wild-type or mutant) octamers wrapped by 5’ biotinylated 147×601 DNA (**Supplemental Table 1)** (Morgan et al. 2021; Marunde et al. 2022). Histone tail truncations were by trypsin digestion of an assembled unmodified nucleosome (tail-less).

This study uses a nomenclature recently devised for accurate scientific communication in the chromatin and epigenetic fields (Keogh et al. 2025). Here ([H2AE61A]_2_) indicates a fully defined homotypic nucleosome where other positions not denoted are understood to be definitively unmodified / wild-type major histones.

### Captify-Alpha binding assays

The assay previously known as dCypher™ is now named Captify™, with no distinction in how the assay is performed or its capabilities. The interaction of Dot5-FLAG or Dot5 (Δ6-11)-FLAG (the Queries: **Supplemental Table 1**) with free DNA (147×601 Widom sequence) or fully defined nucleosomes (the Targets: **Supplemental Table 1**) was assayed by Captify on the Alpha (Amplified luminescence proximity homogeneous assay) platform as previously (Marunde et al. 2022). In brief, queries were serially titrated (in duplicate) against a fixed concentration of target (10 nM biotinylated nucleosome or 2.5 nM free DNA (147×601) and incubated for 30 mins. A 10 μl mix of AlphaScreen streptavidin Donor (*Revvity*, 6760002) and nickel-chelate Acceptor beads (*Revvity*, AL108M) was added, incubated for 60 mins, and Alpha counts measured using a *PerkinElmer* 2104 EnVision plate reader (680 nm laser excitation, 570 nm emission filter ± 50 nm bandwidth). All bead handling steps were in subdued lighting and all incubations at room temperature.

Experiments were performed in 20 mM Tris pH 7.5, 0.01% BSA, 0.01% NP-40, 1 mM DTT with additives as noted (including 50 – 250 mM NaCl and 0 – 20 μg/ml competitor salmon sperm DNA (salDNA)). Binding curves were plotted in GraphPad Prism 9.0 using 4-parameter logistic nonlinear regression to yield EC_50_rel values (**Supplemental Table 3**).

Where necessary, values beyond the Alpha hook point (indicating bead saturation / competition with unbound Query) were excluded and top signal constrained to average max signal for Target. In cases where signal never reached plateau, those were constrained to the average max signal within the assay (relative to unmodified nucleosome). In remaining cases, when a target’s maximal signal never achieved half of max signal relative to unmodified nucleosome, an EC_50_^rel^ was deemed not determinable (ND). In cases where a targets max signal never surpassed a two-fold increase, an EC_50_^rel^ was deemed ND at less than lowest concentration tested (*e.g.*, ND; < 1 nM).

## CUT&RUN

For CUT&RUN, yeast cells expressing Dot5 alleles (**Supplemental Table 1**) were grown and nuclei prepared / stored in aliquots (5 x 10^7^ nuclei per) as previously (Brahma and Henikoff 2022; Aguilar et al. 2025). Nuclei were rapidly thawed at 37 °C, and 100 μL of suspension used per reaction with the CUTANA™ ChIC/CUT&RUN Kit (*EpiCypher* #14-1048). After immunotethering pAG-MNase to Rabbit IgG, anti-H3K4me3 or anti-FLAG (**Supplemental Table 1**), MNase digestion was performed (4 °C for 2 hrs) and DNA eluted in 12 μL final volume. 5 ng of DNA was used to prepare sequencing libraries with the CUTANA CUT&RUN Library Prep Kit (*EpiCypher* #14-1001). Libraries were sequenced on an Illumina NextSeq 2000 platform to a minimum of ∼3 million paired-end (PE) reads per reaction (**Supplemental Table 2**).

PE fastq files were aligned to the *sacCer3* reference genome (Bowtie2(Langmead and Salzberg 2012)), filtered from duplicate (SAMtools (Li et al. 2009)), multi-aligned (broadinstitute.github.io/picard), and exclusion list reads(Lazar-Stefanita et al. 2023), and the resulting unique reads for comparable reactions normalized by a spike-in scaling factor (1/ % *E.coli* reads) (BEDTools v2.30.0 (Quinlan and Hall 2010)) to further RPKM (Reads Per Kilobase per Million mapped reads)-normalized bigwig files (DeepTools). Peak visualization from bigwig files was by Integrative Genomics Viewer (IGV). CUT&RUN sequence data is available from NCBI Gene Expression Omnibus (accession GSE313255). CUT&RUN analyses were performed independently three times with consistent results.

### Structure and disorder predictions

AlphaFold2 (Jumper et al. 2021) was used to generate initial models of each VHH; and AlphaFold 3 (https://alphafoldserver.com/) (Abramson et al. 2024) to predict the structure of Dot5 complexed with a nucleosome. This nucleosome included human histones (also used in our Captify and cryo-EM studies; **Supplemental Table 1**) and 150 bp of randomly generated DNA sequence (147 bp 601 sequence was used in Captify and cryo-EM studies; **Supplemental Table 1**). Contacts between Dot5 and various components of the nucleosome were predicted where distance was < 3 Å. All structures were visualized using Chimera X1.9 (Goddard et al. 2018; Pettersen et al. 2021; Meng et al. 2023).

### Single domain antibodies (VHH) binding to the nucleosome acidic patch

Single domain antibodies (also termed VHH; Variable Heavy Domain of a Heavy chain antibody) to the nucleosome acid patch (chromatibody (Cb) (Jullien et al. 2016); clones 1G1, 1B2 (Hicks et al. 2024)) were created as described. VHH proteins for Captify (VHH-Myc-6xHis: **Supplemental Table 1**) and cryo-EM (VHH-TEV-3xHA-6xHis: **Supplemental Table 1**) studies were expressed as recombinants in *E. coli* (BL21), purified by immobilized metal affinity chromatography, and purity confirmed by SDS-PAGE. VHH binding to the nucleosome acid patch was characterized by Captify-Luminex with the acid patch assessment panel (([H2AE61A]_2_), ([H2AE92K]_2_), ([H2AE105A/E113A]_2_) and unmodified control) as previously (Marunde et al. 2025). Phenotypes resulting from VHH overexpression in yeast (**Supplemental Table 1**) were assayed as in related figure legends.

### VHH-Nucleosome complex formation

VHH (1B2 or 1G1) and unmodified mononucleosome on 5’ biotinylated 147 bp DNA (*EpiCypher* #16-0006) were dialyzed for 3 hrs into complexation buffer (20 mM HEPES pH 7.6, 50 mM NaCl, 0.5 mM MgCl_2_, 1 mM EDTA) in a 3,500 MWCO 0.2 mL dialysis button (*ThermoFisher*) and transferred to 0.5 mL Eppendorf tubes for overnight storage at 4°C. Each VHH was then added at 10:1 (28.8 µL 1B2 at 178 µM or 25.8 µL 1G1 at 199 µM to nucleosome at 10.27 µM) to a total reaction volume of 50 µL. Samples were incubated on ice for 90 minutes, then concentrated using 100 kDa centrifugal filters (*Amicon Ultracel* 0.5 mL) to yield 1.3 mg/mL 1B2-nucleosome and 1.1 mg/mL 1G1-nucleosome. Complex formation was verified on non-denaturing 4-20% Tris-Glycine gels (*BioRad*) and by negative stain electron microscopy with uranyl formate.

### Cryo-EM sample preparation and screening

Grids were prepared and screened at the Penn State Cryo-Electron Microscopy Facility. Quantifoil Cu 200 mesh, R 2/2 grids were glow-discharged for 10 secs at 15 mA in a Pelco easiGlow system. Using an FEI Vitrobot Mk IV maintained at 4 °C with 100% humidity, 3.5 μL of sample was applied to the grid surface, blotted for ∼ 3 secs, and plunged into liquid ethane maintained in liquid nitrogen. Grids were then clipped, loaded onto a Talos Arctica 200 kV equipped with TFS Falcon 4, and best grids selected / images collected for quality control of each sample.

### Cryo-EM Data collection and processing

Datasets were collected at the National Cancer Institute on a Titan Krios 300 kV equipped with a Gatan K3 camera. Super-resolution (81,000x) images (10,279 for 1B2, 8,302 for 1G1) were collected at 1.098 Å / pixel physical size (0.549 Å / pixel super-res). Movies were fractionated into 40 frames at a total dose of 60 e^-^ per Å^2^ (1.25 e^-^ per Å per frame), motion-corrected in Relion (Scheres 2012) using UCSF MotionCor2 (Zheng et al. 2017) and binned from super-resolution 0.549 Å / pixel 2x to 1.098 Å / pixel. These images were then imported to CryoSPARC (Punjani et al. 2017) for further calculations. *“Patch CTF Estimation (multi)”* was used to estimate image defocus, *and “Blob Picker”* to identify particles. Picked particles were extracted in a 300 x 300 box and Fourier-cropped to 80 x 80. 2D classification and selection was then used for each dataset, and best classes selected for “*Ab Initio Reconstruction*”. Data was cleaned up using 2D *“Ab Initio Reconstruction”* and *“Heterogeneous Refinement”*, and resulting best Coulomb potential density maps exported to Relion for *“Bayesian Polishing”* (Zivanov, Nakane and Scheres 2019). Each dataset was then re-imported to CryoSPARC for final refinements. Flowcharts representing data processing are in **Supplemental Figures S10** and **S11.**

Each dataset was calculated independently to avoid biasing. Each dataset faced different challenges: 1B2-Nucleosome was limited by ice thickness, but the VHH bound with high affinity and so could be processed in C2-symmetry. 1G1-Nucleosome had a much better ice thickness but lower occupancy and so was processed in C1-symmetry with focused classification performed in a mask. Final resolutions are reported using the FSC at 0.143 cutoff following gold-standard refinement (Rosenthal and Henderson 2003).

Representative micrographs, 2D classes, local resolution, global resolution and cFSCs are in Supplemental Figure S9.

### Model building

An existing X-ray crystal structure of the nucleosome with 601 Widom sequence (PDB: 3LZ0) (Vasudevan, Chua and Davey 2010) was used as the reference. This was rigid-body fit to each cryo-EM reconstruction, and optimized using UCSF ChimeraX *“Fit in Map”* function (Meng et al. 2023). These nucleosome models were then locally optimized using the *“Real-space refine”* option in Coot (Emsley et al. 2010), and *phenix.real_space_refine* in Phenix (Afonine et al. 2018). Initial models of each VHH were generated using AlphaFold2(Jumper et al. 2021), fit to their respective densities, merged with nucleosome models, manually adjusted, and refined following the same approach as the nucleosome alone. The final refined models were validated automatically using MolProbity, OneDep wwPDB Validation System (Young et al. 2017; Williams et al. 2018), and visually in Coot.

Theoretical models of Dot5 and chromatibody (Cb) in the nucleosomal context were generated using AlphaFold3 (Abramson et al. 2024).

## COMPETING INTEREST STATEMENT

*EpiCypher* is a commercial developer and supplier of reagents (*e.g.*, fully defined semi-synthetic nucleosomes) and platforms (*e.g.*, Captify^TM^ and CUTANA^TM^ CUT&RUN) used in this study. All *EpiCypher* authors are or were employed by (and own shares in) the company, with MWC and MCK also board members of same. The authors declare no additional conflicts.

## ACKNOWLEDGEMENTS

*EpiCypher* is supported by grants R44HG010640, R44GM117683, R44GM136172, R44CA212733 and R43GM134834 from the National Institutes of Health (NIH). JPA is supported by NIH grant R01GM149780. This research was supported in part by the Office of the Director, NIH, under award number S10OD026822-01, and the National Cancer Institute National Cryo-EM Facility at the Frederick National Laboratory for Cancer Research under contract HSSN261200800001E. Cryo-EM data was analyzed on hardware generously provided to the Armache lab through an NVIDIA Academic Grant by NVIDIA Corporation.

We thank Sung Hyun Cho at the Penn State Cryo-Electron Microscopy Facility for assistance with EM screening and data collection, Adam Wier and Thomas Edwards for their support and data collection at the Frederick National Laboratory. JTK is supported by NIH grant R35GM118060. JKT is supported by NIH grant R35GM139816.

## AUTHOR CONTRIBUTIONS

UC constructed yeast strains and performed cell fractionations, phenotypic analyses, structure predictions and immunoblots (with assistance from NA). ECS performed cryo-EM data analysis and interpretation, and prepared structural figures under supervision of JPA. HJF prepared cryo-EM grids. GK identified the HMGN-like motif in Dot5 and purified recombinant Dot5 proteins, while GCB performed EMSAs under supervision of JTK. LFK performed Captify assays under supervision of MRM. RA and MRM performed CUT&RUN with support in data analysis from BJV. SLG, SRH, NLH, KEM and EV produced VHH and dissected their binding to nucleosomes (made by NKS) with support from MWC and ZWS. MCK and JKT provided project conception and supervision, and support in data analysis and interpretation. UC drafted the original manuscript with support from MCK and JKT. All authors contributed to and are responsible for subsequent versions.

## ABBREVIATIONS

Dot: Disruptor of Telomeric Silencing
HMGN: High Mobility Group N
PTM: post-translational modification
salDNA: salmon sperm DNA
VHH: Variable Heavy domain of a Heavy chain.

